# Dual MAPK and HDAC inhibition rewires the apoptotic rheostat to trigger colorectal cancer cell death

**DOI:** 10.1101/2021.02.15.431252

**Authors:** Laura J. Jenkins, Ian Y. Luk, Michelle Palmieri, W. Douglas Fairlie, Erinna F. Lee, Kael L. Schoffer, Tao Tan, Irvin Ng, Natalia Vukelic, Janson W.T. Tse, Rebecca Nightingale, Zakia Alam, George Iatropoulos, Matthias Ernst, Shoukat Afshar-Sterle, Jayesh Desai, Peter Gibbs, Oliver M. Sieber, Amardeep S. Dhillon, Niall C. Tebbutt, John M. Mariadason

**Affiliations:** Olivia Newton-John Cancer Research Institute, Melbourne, Vic., Australia; School of Cancer Medicine, La Trobe University, Melbourne, Vic., Australia; Department of Biochemistry and Genetics, La Trobe Institute for Molecular Science, La Trobe University, Melbourne, Vic., Australia; Personalised Oncology Division, The Walter and Eliza Hall Institute of Medical Research, Melbourne, Vic., Australia; Peter MacCallum Cancer Centre, Melbourne, Vic., Australia; Department of Medical Biology, The University of Melbourne, Melbourne, Vic., Australia; Department of Surgery, The University of Melbourne, Melbourne, Vic., Australia; Department of Biochemistry and Molecular Biology, Monash University, Melbourne, Vic., Australia; The Institute for Mental and Physical Health and Clinical Translation, School of Medicine, Deakin University, Geelong., Australia

**Keywords:** Colon cancer, MAPK, HDAC inhibitor, apoptosis

## Abstract

The EGFR/RAS/MEK/ERK signalling pathway (ERK/MAPK) is hyper-activated in most colorectal cancers (CRCs). A current limitation of inhibitors of this pathway is that they primarily induce cytostatic effects in CRC cells. Nevertheless, these drugs do induce expression of pro-apoptotic factors, suggesting they may prime CRC cells to undergo apoptosis. As histone deacetylase inhibitors (HDACi) induce expression of multiple pro-apoptotic proteins, we examined whether they could synergize with ERK/MAPK inhibitors to trigger CRC cell apoptosis. Combined MEK/ERK and HDAC inhibition synergistically induced apoptosis in CRC cell lines and patient-derived tumour organoids *in vitro,* and attenuated *Apc*-initiated adenoma formation *in vivo*. Mechanistically this effect was mediated through induction of the BH3-only pro-apoptotic proteins BIM and BMF. Importantly, we demonstrate that this treatment paradigm can be tailored to specific MAPK genotypes in CRCs, by combining HDACi with EGFR, KRAS^G12C^ or BRAF^V600^ inhibitors in *KRAS/BRAF*^*WT*^; *KRAS*^*G12C*^, *BRAF*^*V600E*^ CRC cell lines respectively. These findings identify a novel ERK/MAPK genotype-targeted treatment paradigm for colorectal cancer.

## Introduction

The ERK/MAPK signalling pathway is constitutively activated in a high percentage of colorectal cancers (CRC). This is driven by either activating mutations in *KRAS, NRAS* or *BRAF* which collectively occur in ~55% of cases [1], or as a result of upregulated canonical activation of receptor tyrosine kinases such as EGFR [2].

Activation of ERK/MAPK signalling in cancer cells drives cell proliferation, and is associated with loss of cell differentiation and promotion of cell survival [3]. The pro-survival effects of the pathway are illustrated by the transcriptional repression of pro-apoptotic BH3-only proteins such as BIM [4] and PUMA [5]. ERK-mediated phosphorylation of BIM_EL_ also promotes its ubiquitination and degradation [6–8], as well as its dissociation from the pro-survival proteins BCL-X_L_, BCL-2 and MCL-1 [8]. ERK also phosphorylates BMF, which reduces BMF-dependent cell-killing activity [9], and induces RSK-mediated phosphorylation of BAD, which sequesters BAD to the cytosol [10, 11]. Finally, ERK signalling promotes expression of BCL-2 like pro-survival proteins. For example, in *BRAF* mutant tumours ERK phosphorylates MCL-1 on Thr163 promoting its stabilisation [12]. Conversely, ERK/MAPK inhibitors (MAPKi) have been shown to induce the opposite effect in cancer cell lines, increasing expression of BIM and BMF, and downregulating MCL1 [13–15].

Despite these effects, ERK/MAPK inhibitors (ERK/MAPKi) primarily induce cytostatic effects in tumour cells, including CRC cells [16]. This is also supported by clinical observations where liquid biopsy monitoring studies of patients treated with EGFR inhibitors revealed that levels of mutant RAS clones increase in the blood during EGFR blockade, but then decline following drug treatment withdrawal, indicating preferential regrowth of the original dominant clones [17, 18]. Collectively these findings suggest that ERK/MAPKi may be unable to alter the apoptotic “rheostat” to sufficient levels needed to trigger apoptosis, but may induce a “primed” apoptotic state in tumour cells, which may be exploited through combination drug treatment.

Histone deacetylase inhibitors (HDACi) are a class of epigenetic modulators approved for treatment of the haematological malignancies cutaneous T-cell lymphoma and multiple myeloma. A number of HDACi are in routine clinical use including vorinostat (Zolinza, Merck), panobinostat (Farydak, Novartis) and romidepsin (Istodax, Celgene). While HDACi induce multiple effects on tumour cells, pre-clinical studies indicate that their anti-tumour activity is predominantly driven through the induction of apoptosis [19]. Depending on the cell type, these pro-apoptotic effects are triggered, at least in part, through the transcriptional activation of pro-apoptotic mediators including BAD [20], BIK [20], NOXA [20], BMF [21–23], BIM [24] and BAK [25] as well as the repression of pro-survival proteins including BCL-X_L_ [20, 26], BFL-1 [20], BCL-w [20] and BCL-2 [27].

While HDACi have shown limited single agent activity in solid tumours, their unique mechanism of action makes them well suited for use in rational combination regimens and represent a potential class of drugs that may enhance that apoptotic activity of MAPK pathway inhibitors. To address this possibility, we investigated if HDACi enhances the apoptotic activity of MAPKi in CRC cells, and whether this paradigm could be tailored to target CRCs with different MAPK pathway lesions. We also investigated the mechanism of action of this therapeutic combination in relation to altering the apoptotic rheostat in CRC cells.

## Materials and methods

### Cell culture

CRC cell lines used in this study were obtained from the American Type Culture Collective (Manassas, VA, USA) or other investigators as previously described [28, 29]. All cell lines were maintained in Dulbecco’s Minimal Essential Media/F12 (DMEM/F12) supplemented with 10% FBS (v/v) (all from Thermo Fisher Scientific, Waltham, MA, USA) at 37°C with 5% CO_2_.

### Chemicals

Trametinib, panobinostat, vorinostat, mocetinostat, entinostat, romidepsin and vemurafenib (Selleck Chemicals, Houston, TX, USA) and AMG 510 (Active Biochem, Kowloon, Hong Kong) were dissolved in dimethyl sulfoxide (DMSO, Sigma-Aldrich, St. Louis, MO, USA). Valproic acid (Sigma-Aldrich) and dissolved in water. Cetuximab (Erbitux, Lilly, Indianapolis, IN, USA) was obtained from the Austin Health Pharmacy.

### Cell viability

Cells were seeded in 96 well plates at densities optimised to ensure control cells did not reach confluence during the experimental period. Cell viability was determined using the MTS assay (Promega, Madison, WI, USA), by incubating cells for 90 minutes at 37°C and measuring absorbance at 490 nm (MTS) and 630 nm (background absorbance) on a SPECTROstar Nano Spectrophotometer (BMG LabTech, Germany).

### Flow cytometry

Cells were seeded in 24 well plates and treated with drug for 24-72 hours. Adherent and non-adherent cells were then collected and incubated in 50 μg/mL propidium iodide (Sigma-Aldrich), 0.1% sodium citrate (w/v) and 0.1% Triton X-100 (v/v) (Sigma-Aldrich) overnight at 4°C. Samples were then analysed on a FACS Canto II flow cytometer (BD Biosciences, San Jose, CA, USA), by analysis of 10,000 events. Apoptotic cells were defined as those with a sub-diploid content and quantified using FlowJo V8.0 (FlowJo LLC, Ashland, OR, USA).

### Patient-derived tumour organoids (PDTOs)

PDTO’s were generated from resected tumour material from de-identified patients with colorectal cancer. All patients provided informed consent and the study was approved by a multi-institutional Human Research Ethics Committee (HREC 2016.249). Organoids were resuspended in DMEM/F-12 with Matrigel (Corning, NY, USA) as a single cell suspension and seeded at 3,000 cells per well into a 384 optical well plate (Nunc, Denmark), and allowed to establish for 72 hours before the addition of drug. Drug combinations were prepared with 5 μg/mL of propidium iodide (Sigma-aldrich) in a matrix formation using panobinostat (1, 5, 10, 25 and 50 nM) and trametinib (1, 3, 5, 10 and 25 nM). Organoids were imaged on a Nikon Ti2 Eclipse fluorescent microscope (Nikon, Tokyo, Japan) at 4x magnification at a constant temperature and CO_2_ levels throughout the duration of the experiment. Images were acquired at 72 hours, with a z-stack of 21 slices at 25 μm intervals. At endpoint, viability was determined using the CellTiter-Glo 3D (Promega) viability assay and luminescence read on the EnVision Plate Reader (PerkinElmer, Waltham, MA, USA). Fluorescence activity data acquired from the Envision Plate Reader was transformed into relative activity, with DMSO as negative (vehicle) control and 1 μM Bortezomib as a positive (killing) control.

### *Apc*^*lox/lox*^;*Cdx2-Cre*^*ERT2*^ Mice

*Apc*^*lox/lox*^;*Cdx2-Cre*^*ERT2*^ mice were generated as originally described [30]. Cre-recombinase mediated *Apc* gene inactivation was induced in 5 week old mice by intraperitoneal injection of 1 mg (100 μL of 10 mg/mL solution) of tamoxifen for two consecutive days. Mice were then left for 1 week to allow for adenoma initiation after which drug treatment was commenced. Mice were then randomised into 4 groups to receive either; vehicle control, trametinib (0.3 mg/kg, oral gavage), panobinostat (5 mg/kg, IP injection) or trametinib 0.3 mg/kg and panobinostat 5 mg/kg. Mice were treated with drug for five consecutive days followed by 2 days of rest and received a total of 12 treatments. The study was approved by the Austin Health Animal Ethics Committee.

### Endoscopy procedure

Mice were anaesthetised using (1 L O_2_/min and 2-3 % Isoflurane). The procedure was performed using a miniature endoscope (scope 1.9 mm outer diameter), a light source, an IMAGE 1 camera, and an air pump (all from Karl Storz, Germany), to achieve regulated inflation of the mouse colon.

### Xenograft studies

For xenograft studies, 6 weeks old female BALB/c-Foxn1^nu^/ARC (BALB/c nude) mice were purchased from the Animal Resources Centre (Western Australia, Australia) and housed in specific pathogen free (SPF) microisolators. Mice were then subcutaneously injected with 2 million LIM1215 cells into the left and right flanks in a 1:1 mixture of matrigel matrix (75 μL)(Corning): DMEM-F12 (75 μL). Once tumours became palpable, mice were randomised to receive either; vehicle control, cetuximab (40 mg/kg, IP injection), panobinostat (5 mg/kg, IP injection) or cetuximab 40 mg/kg and panobinostat 5 mg/kg, for five consecutive days for a total of 10 treatments. The study was approved by the Austin Health Animal Ethics Committee.

### Immunohistochemistry

Formalin fixed paraffin embedded (FFPE) sections were de-paraffinised and rehydrated through serial washes in xylene and ethanol. Sections were rinsed in H_2_O and quenched in 3% H_2_O_2_ (Chem -supply, SA, Australia) for 10 minutes. Antigen retrieval was performed by incubation in Citrate buffer (pH 6.0) in a boiling water bath for 30 minutes. Slides were probed with β-catenin (C19220, BD Transduction Laboratories, San Jose, CA, USA) and phosho-p44/42 MAPK (ERK1/2) Thr202/Thr204 (4370, Cell Signalling Technologies, Danvers, MA, USA) at 4°C overnight, then washed and incubated with Labelled polymer HRP-anti-Rabbit and Labelled polymer HRP-anti-mouse secondary antibody (Dako, Denmark) for 1 hour at room temperature. Chromagen was developed using the DAB (3, 3-diaminobenzidine) reagent (Dako). Sections were counter-stained using pre-filtered Mayer’s haematoxylin (Amber Scientific, Australia) then dehydrated through serial ethanol and xylene washes before mounting using DPX mounting solution (Sigma-aldrich).

### Western blot analysis

Protein isolation was performed using NP-40 lysis buffer (50 mM Tris-HCl pH 7.5, 150 mM NaCl, 1% NP-40, 0.5% sodium deoxycholate, 1 mM EDTA pH 8 and 1 tablet of protease inhibitor cocktail Roche cOmplete (Roche, Basel, Switzerland) and 1 tablet of phosphatase inhibitor PhosSTOP (Roche)). A total of 30 μg of protein per sample was separated on NuPAGE 4-12% Bis-Tris pre-cast polyacrylamide gels (Novex, Thermo Fisher Scientific), transferred onto iBLOT2 Polyvinylidene Difluoride (PVDF) membranes (Invitrogen, Thermo Fisher Scientific) then blocked using Odyssey blocking buffer (LiCor, Lincoln, NE, USA). Primary antibodies used in western blot analysis were BIM_S/EL/L_ (ALX-804-527, Enzo Life Sciences, New York, NY, USA), BMF (ALX-804-343-C100, Enzo Life Sciences), MCL-1 (5453S, Cell Signalling Technologies), phosho-p44/42 MAPK (ERK1/2) Thr202/Thr204 (4370, Cell Signalling Technologies), p44/42 MAPK (ERK1/2) (9107, Cell Signalling Technologies), β-actin (A5316, Sigma-aldrich) and β-tubulin (ab6046, Abcam, Cambridge, UK). Secondary antibodies used include, IRDye®680RD Goat anti-mouse IgG (H+L), IRDye®800CW Goat anti-rabbit IgG (H+L) and IRDye®680RD Goat anti-rat IgG (H+L) (LiCor). Signal was detected using an Odyssey imaging system (LiCor)

### Quantitative-Real Time-PCR (q-RT-PCR)

RNA was isolated using the ReliaPrep™ RNA tissue miniprep system (Promega) as per the manufacturer’s protocol. cDNA was synthesised from 1 μg of RNA using the High-Capacity cDNA Reverse Transcription Kit (Applied Biosystems, Thermo Fisher Scientific) as per the manufacturers protocol. Quantitative RT-PCR was performed using SYBR green master mix (Applied Biosystems) on the Viia™ 7 Real-Time PCR system. A list of primers used is listed in **Table S1**.

### siRNA-mediated knockdown

Knockdown experiments were performed by transfecting cells when 60% confluent with a mixture of Lipofectamine RNAiMax (Invitrogen, Thermo Fisher Scientific) and 25 nM Non-Targeting (NT) or BMF-targeting siRNAs (Bioneer, Daejeon, South Korea). Cells were transfected for 24 hours prior to drug treatment. BMF targeting siRNAs comprised of the following sequences siBMF#1 (5’-UAGUGAAGCUGCUAUCCU) and siBMF#2 (5’-UCUAAAAGUCACUUAGCU).

### Generation of BIM knockout cells using CRISPR-Cas9

BIM knockout CRC cell lines were generated using a previously described doxycycline inducible CRISPR-Cas9 lentiviral platform with the following sgRNA sequence: *hBim* exon 3: 5’-GCCCAAGAGTTGCGGCGTAT [31]. Independent lentiviruses expressing Cas9-mCherry and hBIM-guide RNA-GFP were generated by transfecting lentiviral components into 293T packaging cells, and transduced into COLO 201 CRC cells. GPF/mCherry double-positive cells were subsequently sorted using a BD FACSAria™ II (BD Biosciences). Once expanded, expression of the BIM sgRNA was induced by treatment with 1 μg doxycycline-hyclate (Sigma-Aldrich), and BIM knockout confirmed by western blot.

### Statistical analysis

Statistical analysis was performed using GraphPad Prism v8.0 software (GraphPad Software, La Jolla, CA, USA) whereby experiments containing more than two groups were analysed using One-Way ANOVA, with Tukey’s multiple comparison testing. Experiments containing comparison between two groups were analysed using the student’s T-test with Welch’s correction. In all cases *P*<0.05 was considered statistically significant. Synergy was analysed using two separate models; Bliss Excess and the Highest Single Agent (HSA). All calculations and specific functions were performed in R Version 3.0.1 (R Core Team (2012)). R: A language and environment for statistical computing. R Foundation for Statistical Computing, Vienna, Austria. ISBN 3-900051-07-0, URL http://www.R-project.org/).

## Results

### Combined MEK and HDAC inhibition synergistically induces apoptosis in CRC cell lines with different MAPK pathway lesions and in patient-derived colorectal tumour organoids

To assess the potential of combinatorial inhibition of ERK/MAPK signalling and HDAC activity as a therapeutic strategy in CRC, we treated nine CRC cell lines harbouring different MAPK pathway genotypes with the MEK inhibitor (MEKi) trametinib and the HDACi panobinostat for 72 hours. The combination treatment induced significantly higher apoptosis compared to either agent alone in all cell lines (**Figure 1**), including *BRAF* mutant lines (**Figure 1A**), *KRAS* mutant lines (**Figure 1B**), and *KRAS/BRAF* wildtype lines including EGFR-amplified DiFi cells (**Figure 1C**).

**Figure 1.**
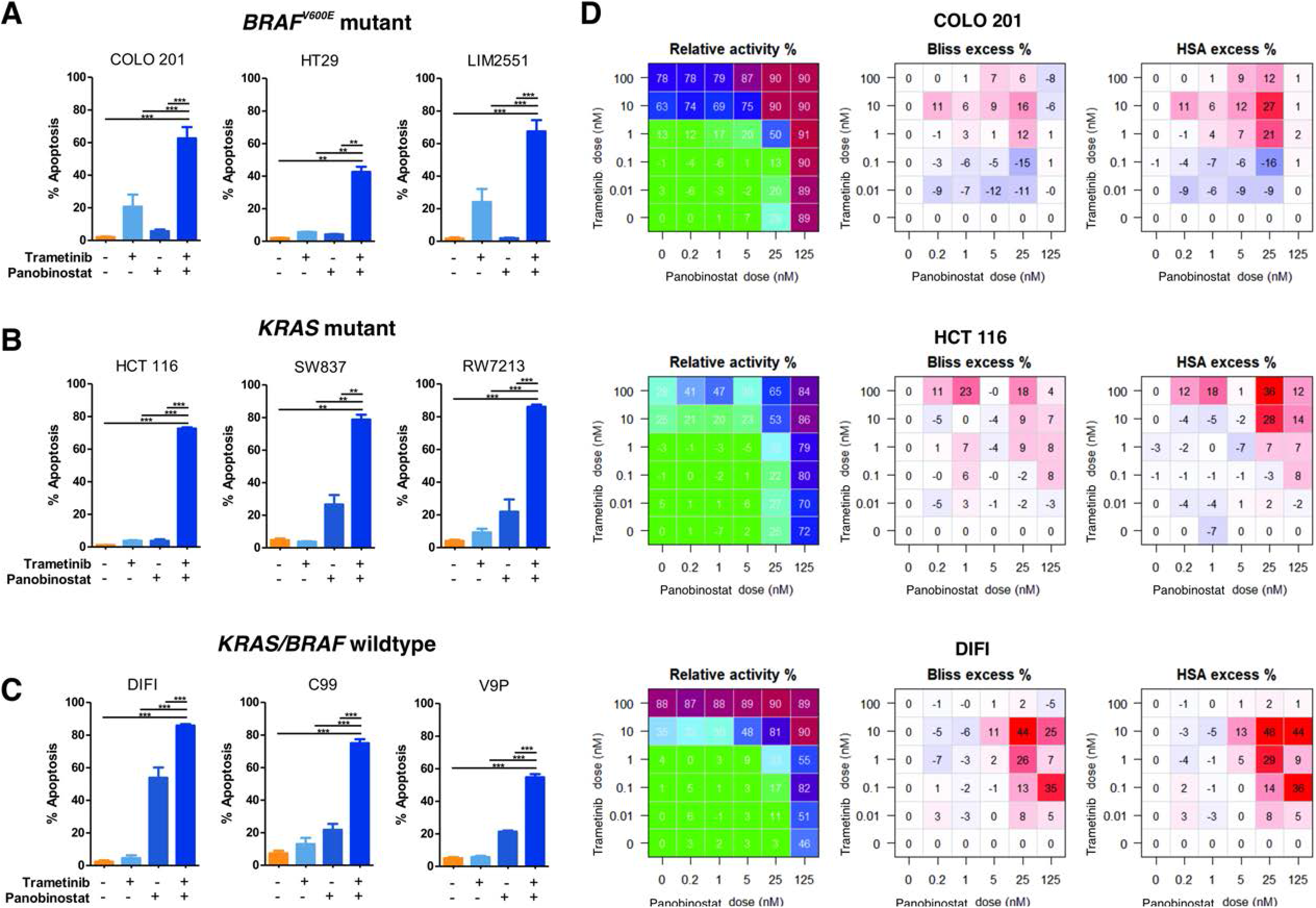
Combined MEK/HDAC inhibition induces apoptosis and reduces viability of CRC cells. (**A**) *BRAF*^*V600E*^ mutant, (**B**) *KRAS* mutant and (**C**) *KRAS*/*BRAF* wildtype CRC cell lines were treated with trametinib (10 nM) and panobinostat (25 nM) alone or in combination for 72 hours and apoptosis induction determined by propidium iodide staining and FACS analysis. Values shown are mean ± SEM from three independent experiments. Differences were compared using an One-Way ANOVA, with Tukey’s multiple comparison testing; **(*p*≤0.01) and *** (*p*≤0.001). (**D**) Effect of trametinib and panobinostat combination on cell viability. *BRAF*^*V600E*^ mutant (COLO 201), *KRAS* mutant (HCT 116) and *BRAF*/*KRAS* WT (DIFI) CRC cell lines were treated in a matrix of increasing concentrations of trametinib and panobinostat for 72 hours and cell viability assessed using the MTS assay. Data shown are from a single representative experiment performed in quadruplicate. Synergy, additivity or antagonism was assessed by using the Bliss excess percentage and HSA excess percentage methods. Synergy is denoted by darker red colours whereas blue is indicative of antagonism.

To assess if the effects of the MEKi/HDACi combination was synergistic, three cell lines with different MAPK pathway genotypes were treated with a range of concentrations of trametinib and panobinostat, and synergy assessed following MTS assays by Bliss and HSA excess models. Synergistic activity was confirmed in all 3 cell lines (**Figure 1D**).

To explore the mechanism of MEKi/HDACi-induced cell death, HT29 cells were treated with the drug combination in the presence of the pan-caspase inhibitor Q-VD-OPh. Cell death was significantly attenuated by Q-VD-OPh, indicating that apoptosis induction was caspase-dependent (**Figure S1**).

As panobinostat belongs to the hydroxamic acid class of HDACi, we next assessed if the synergistic apoptotic activity of the MEKi/HDACi combination extended to other structural classes of HDACi. As with panobinostat, combining trametinib with another clinically used hydroxamic acid vorinostat, the cyclic tetrapeptide romidepsin, the benzamides mocetinostat or entinostat, or the short-chain fatty acid valproic acid, all significantly enhanced apoptosis compared to either agent alone (**Figure S2**).

To confirm these findings in an independent model system, we tested the effects of this combination in four patient-derived colorectal tumour organoids (PDTO’s), three of which were derived from metastatic lesions and one from a stage II tumour. As observed in CRC cell lines, the MEKi/HDACi combination synergistically induced cell death in all four tumour organoid lines as measured by MTT assay, and confirmed by the uptake of Propidium Iodide and membrane blebbing characteristic of apoptosis (**Figure 2**).

**Figure 2.**
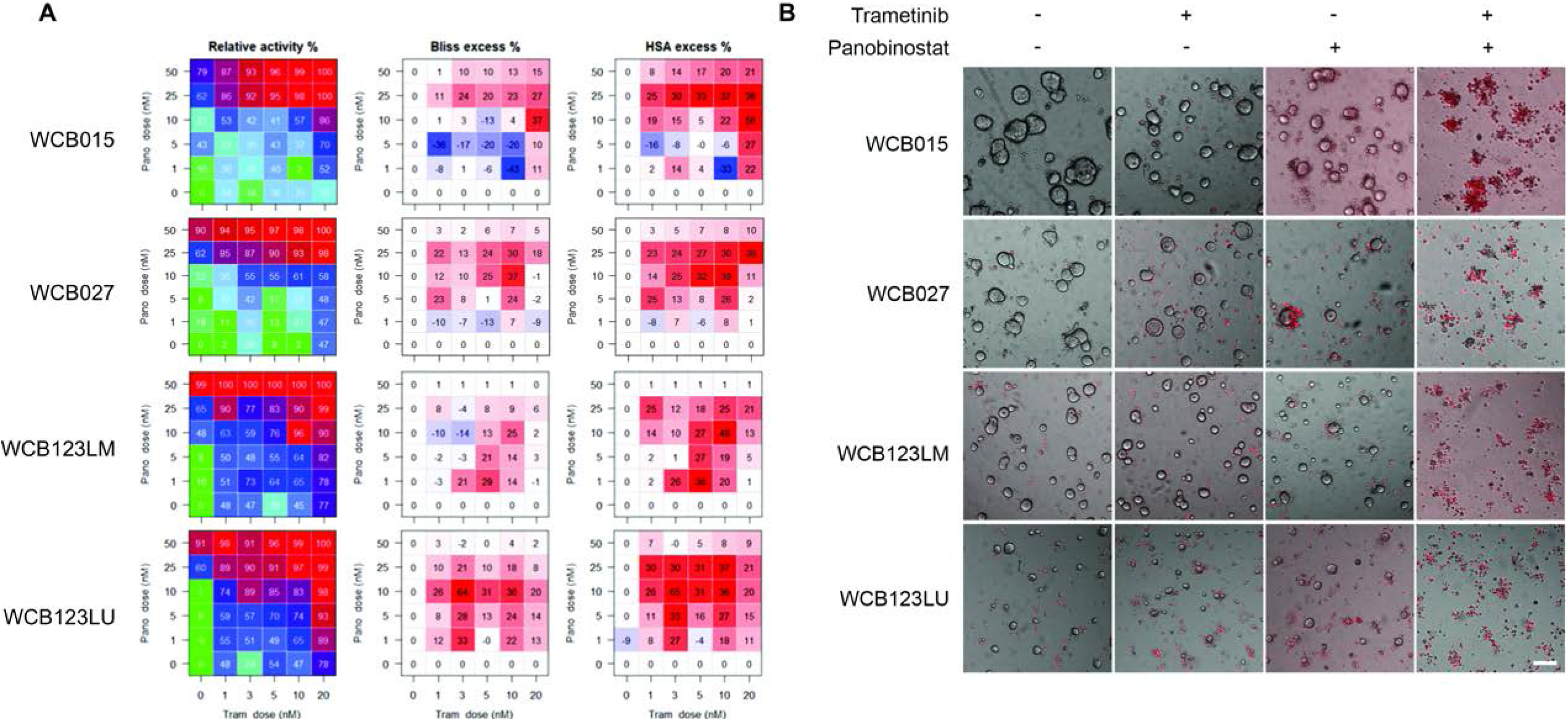
Combined MEK/HDAC inhibition reduces viability of patient-derived colorectal tumour organoids. (**A**) Assessment of synergistic drug activity in PDTO’s WCB015, WCB027, WCB123LM and WCB123LU treated in a matrix of trametinib and panobinostat for 72 hours. Synergy, additivity or antagonism was assessed by using the Bliss excess percentage and HSA excess percentage methods. Synergy is denoted by darker red colours whereas blue is indicative of antagonism. (**B**) Representative images of propidium iodide stained PDTO’s treated with trametinib (10 nM) and panobinostat (25 nM) alone and in combination for 72 hours. Images were obtained at 4x magnification. Cells which have incorporated propidium iodide are stained red.

### MEK/HDAC co-inhibition attenuates *Apc*-initiated colon adenoma development

To assess the efficacy of combined MEK/HDAC inhibition in earlier stage disease, we tested the drug combination in *Apc*^lox/lox^;*Cdx2*-*Cre*^*ERT2*^ mice, which develop multiple colonic adenomas 30 days after tamoxifen-induced inactivation of the *Apc* tumour suppressor gene. As expected, immunohistochemical (IHC) staining of colonic adenomas confirmed high nuclear expression of β-catenin, consistent with activation of Wnt signalling. Importantly, IHC staining for pERK revealed that the ERK/MAPK pathway was also robustly induced in these adenomas (**Figure S3**). To test the activity of MEK/HDAC inhibition in this model, tamoxifen-induced *Apc*^*lox/lox*^;*Cdx2-Cre*^*ERT2*^ mice were treated with vehicle, trametinib, panobinostat or the combination for two weeks. Endoscopic assessment of adenoma development on the day following the final drug treatment revealed fewer adenomas in mice treated with trametinib or panobinostat alone compared to control, and a further reduction in mice treated with the combination (**Figure 3A**). To quantify this effect, an unbiased macroscopic count of adenomas was performed at cull which confirmed that tumour number and size was lowest in mice treated with the MEKi/HDACi combination (**Figure 3B-D**). Importantly, while single agent treatments resulted in an ~10% loss of overall body weight, the combination did not exacerbate these effects (**Figure S4A**)

**Figure 3.**
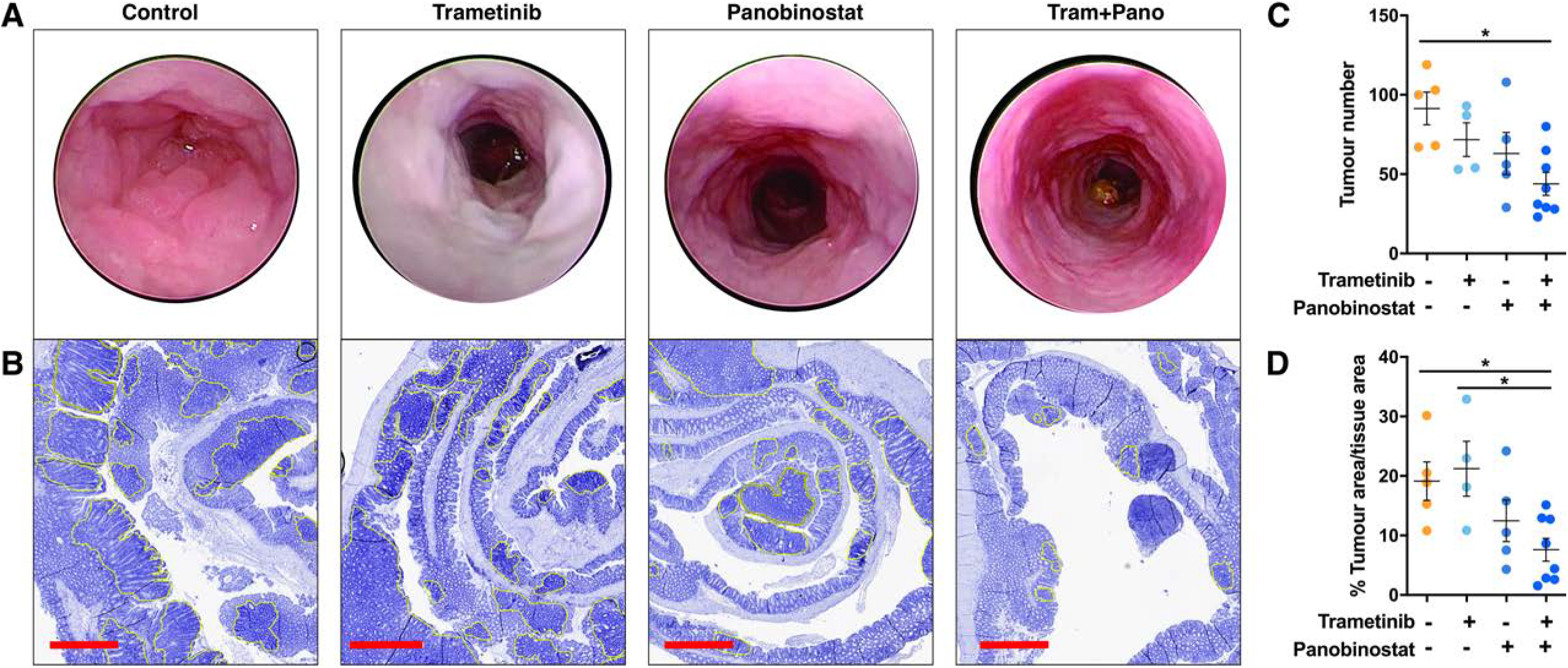
Combined MEK/HDAC inhibition attenuates colonic adenoma formation. (**A**) Representative endoscopy images of control and drug treated mice obtained immediately prior to the experimental endpoint. Mice were treated with trametinib (0.3 mg/kg), panobinostat (5 mg/kg) or the combination for five consecutive days followed by 2 days of rest and received a total of 12 treatments. (**B**) Representative H&E stains of the entire colon from mice treated with vehicle control, trametinib, panobinostat or trametinib plus panobinostat. Yellow outlining indicates tumours. Scale bar = 900 μm. (**C**) Tumour number obtained from macroscopic counting of adenomas in resected colons. (**D**) Tumour burden as determined by computing total tumour area relative to total tissue area from haematoxylin and eosin (H&E) stained sections of the entire colon, from vehicle control (n=5), trametinib (n=4), panobinostat (n=5) and combination (n=8) treated mice. Values shown are mean ± SEM from the biological replicates. One-Way ANOVA, with Tukey’s multiple comparison testing; *(*p*≤0.05), *** (*p*≤0.001).

### Combined MEK/HDACi inhibition additively induces expression of BH3-only genes

MAPKi and HDACi have been independently shown to alter expression of multiple components of the intrinsic apoptotic pathway [20, 32]. We therefore sought to identify pro- and anti-apoptotic genes whose expression was altered by combined MAPK/HDAC inhibition across multiple CRC cells, with different MAPK pathway genotypes (**Figure S5**). While expression of several apoptotic regulators was altered in individual cell lines, the only factor we found to be consistently induced across all four cell lines was the BH3-only gene *BMF* (**Figure S5**), which was further confirmed at the protein level (**Figure 4A**).

**Figure 4.**
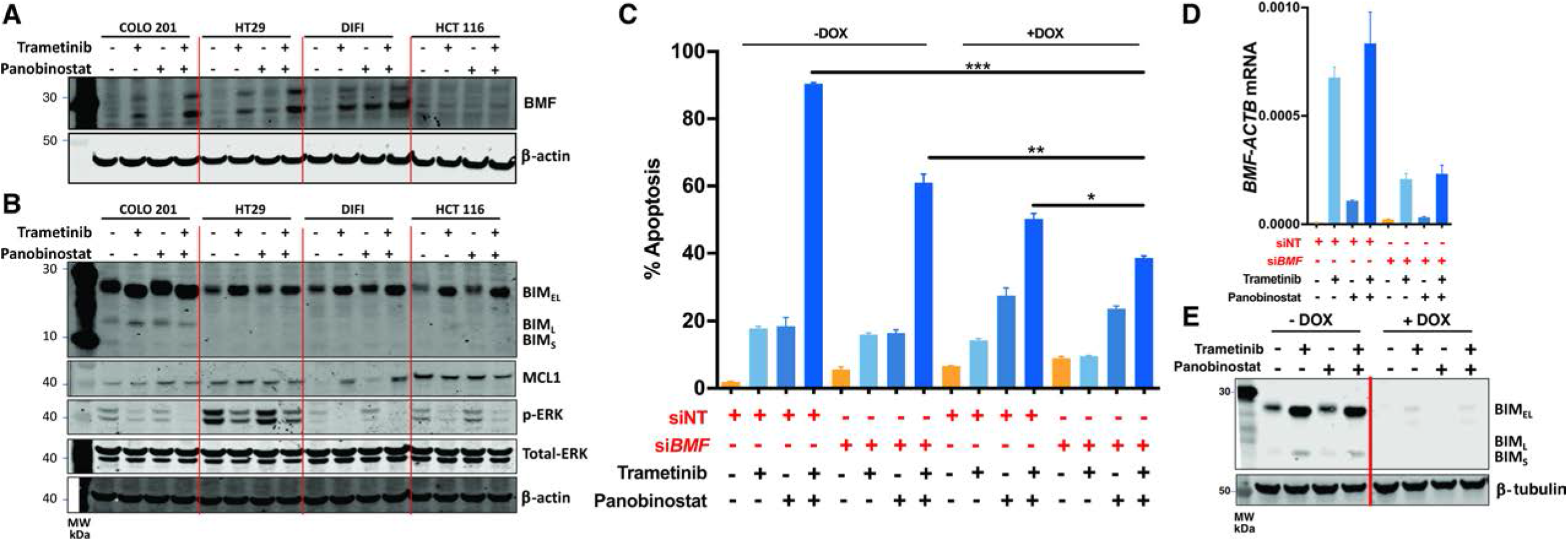
Induction of BMF and BIM is required for MEK/HDACi-induced apoptosis. Effect of trametinib (10 nM) and panobinostat (25 nM) treatment in COLO 201, HT29, DIFI and HCT 116 on (**A**) BMF protein expression following 24 hours treatment and (**B**) BIM, MCL-1, pERK and total-ERK following 6 hours of treatment as determined by western blot. β-actin was used as a loading control. (**C**) Effect of dual inactivation of BIM and BMF on trametinib and panobinostat induced apoptosis. COLO 201 BIM deletion CRISPR cells (minus doxycycline (−DOX) and plus doxycycline (+DOX)) were transiently transfected with non-targeting or BMF-targeting siRNAs for 24 hours. Cells were then treated with trametinib (10 nM) and panobinostat (25 nM) alone and in combination for 24 hours and apoptosis determined by propidium iodide staining and FACS analysis. (**D**) Corresponding assessment of the knockdown efficiency of *BMF* by q-RT-PCR. (**E**) Validation of CRISPR-Cas9-mediated BIM deletion by western blot. β-tubulin was used as a loading control. Values shown are mean ± SEM from a single experiment performed in technical triplicate. Similar results were obtained in a second independent experiment. One-Way ANOVA, with Tukey’s multiple comparison testing; *(*p*≤0.05), **(*p*≤0.01) and ***(*p*≤0.001)

MAPK/ERK signalling also results in the phosphorylation and degradation of BIM_EL_ [6-8, 33] and stabilisation of MCL-1 [12]. We therefore also examined expression of these proteins following combined MEK/HDAC inhibition in the same cell lines. MEKi alone increased BIM expression as well as an increase in the mobility of BIM_EL_ (or decrease in the apparent molecular mass) consistent with its expected dephosphorylation following MAPK/ERK pathway inhibition, while HDACi treatment induced a more modest increase in BMF expression (**Figure 4B**). Comparatively, expression of MCL-1 expression was not consistently changed across the four cell lines following either single agent or combination treatment (**Figure 4B**). As expected, p-ERK levels were decreased in cells treated with trametinib alone, or in combination with panobinostat, while panobinostat alone had no effect (**Figure 4B**). Collectively, these findings demonstrate that combined MEK/HDAC inhibition consistently increased expression of BMF and BIM across multiple CRC cell lines of different MAPK pathway genotypes.

### MEKi/HDACi-induced apoptosis in CRC cells is dependent on BIM and BMF

To determine whether the induction of BMF and BIM is required for MEKi/HDACi-induced apoptosis, we first deleted BIM in COLO 201 CRC cells using CRISPR/Cas9 (**Figure 4C-E**). BMF was subsequently downregulated in BIM knockout cells by transient transfection with BMF-targeting siRNAs. Efficient inactivation of BMF and BIM under basal conditions or following MEKi/HDACi treatment was confirmed by q-RT-PCR and western blot respectively (**Figure 4D-E**). While inactivation of either BMF or BIM alone resulted in partial attenuation of MEKi/HDACi-induced apoptosis, combined inactivation of both factors resulted in a significantly greater attenuation in apoptosis (**Figure 4C**), demonstrating BMF and BIM are both required for apoptosis induction by MEK/HDAC inhibition.

### Tailoring combined MAPK pathway and HDAC inhibition to the MAPK pathway mutation status of colorectal cancer

MAPK/ERK signalling is constitutively activated in CRC by multiple mechanisms including canonical EGFR-driven signalling, *RAS* and *BRAF* mutations. Multiple therapies have been developed to selectively inhibit the pathway at specific nodes. These include EGFR-inhibitors, BRAF inhibitors and most recently G12C mutant KRAS inhibitors. We therefore sought to determine the efficacy of combining HDACi with a specific MAPKi tailored to the MAPK pathway genotype of the tumour.

To address this, we first tested the effect of combining panobinostat with the EGFR-targeting antibody (cetuximab) in six *KRAS*/*BRAF* wildtype CRC cell lines. Combined EGFR/HDAC inhibition significantly enhanced apoptosis compared to treatment with vehicle or either agent alone in all cell lines (**Figure 5A**). To validate this finding *in vivo*, we tested the efficacy of combining cetuximab and panobinostat on growth of LIM1215 xenografts. While both cetuximab and panobinostat alone significantly reduced tumour volume compared to vehicle control, a further reduction was observed in mice treated with the drug combination (**Figure 5B**). This effect was confirmed when the weight of resected tumours was assessed (**Figure 5C**). Importantly, the cetuximab/panobinostat combination was well tolerated by mice as indicated by stable maintenance of body weight for the duration of the experiment in all treatment groups (**Figure S4B**). Similar to the effects induced by MEK/HDAC inhibition, EGFR/HDAC inhibition additively induced *BMF* mRNA and BIM protein expression (**Figure 5D-E**).

**Figure 5.**
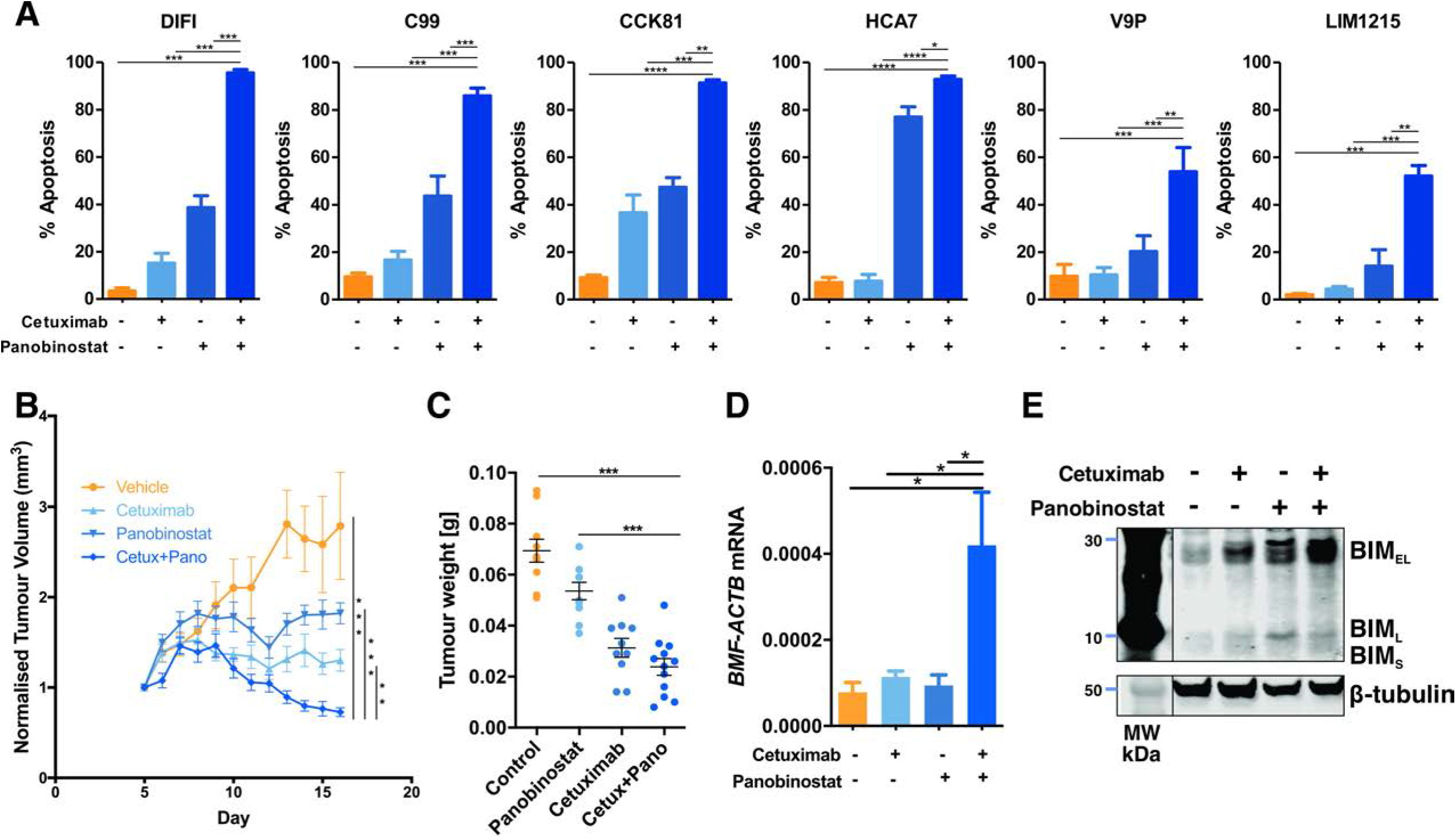
Combined EGFR/HDAC inhibition induces apoptosis in CRC cells. (**A**) *KRAS*/*BRAF* wildtype CRC cell lines treated with cetuximab (10 μg/mL) and panobinostat (25 nM) alone or in combination for 72 hours and apoptosis induction determined by propidium iodide staining and FACS analysis. Values shown are mean ± SEM from three independent experiments. One-Way ANOVA, with Tukey’s multiple comparison testing; *(*p*≤0.05), **(*p*≤0.01) and ***(*p*≤0.001). (**B**) Effect of cetuximab and panobinostat on growth of LIM1215 tumour xenografts. BALB/*c* nude mice per group were subcutaneously injected with 2 x 10^6^ LIM1215 cells into the right and left flanks (n=10-12). Mice were then randomized to receive either vehicle control, cetuximab (n=5, 40 mg/kg), panobinostat (n=5 5 mg/kg) or the combination (n=6) for a total of 12 treatments. Tumour volume was normalised to day 1 of treatment. Values shown are mean ± SEM. (**C**) Weight (g) of excised tumours at experimental endpoint. (**D**) Expression of *BMF* mRNA expression in excised tumours (n=3, all groups). Values shown are mean ± SEM. One-Way ANOVA, with Tukey’s multiple comparison testing; *(*p*≤0.05), **(*p*≤0.01), *** (*p*≤0.001). (**E**) Expression of BIM in representative excised tumours from each treatment group as determined by western blot analysis. β-tubulin was used as a loading control.

We next assessed the effect of combining panobinostat with the KRAS^G12C^ inhibitor AMG 510 in *KRAS*^*G12C*^ mutant cell lines. The selectivity of AMG510 for KRAS^G12C^ was first confirmed by demonstrating its ability to selectively inhibit cell proliferation and ERK/MAPK signalling in *KRAS*^*G12C*^ mutant CRC cells (**Figure 6A-B**). As observed for the previous MAPKi/HDACi combinations, KRAS^G12C^/HDAC inhibition significantly increased apoptosis induction compared to either agent alone in two *KRAS*^*G12C*^ cell lines (**Figure 6C**), and further increased expression of BMF (**Figure 6E, G**) and BIM, albeit to a lesser extent (**Figure 6G**).

**Figure 6.**
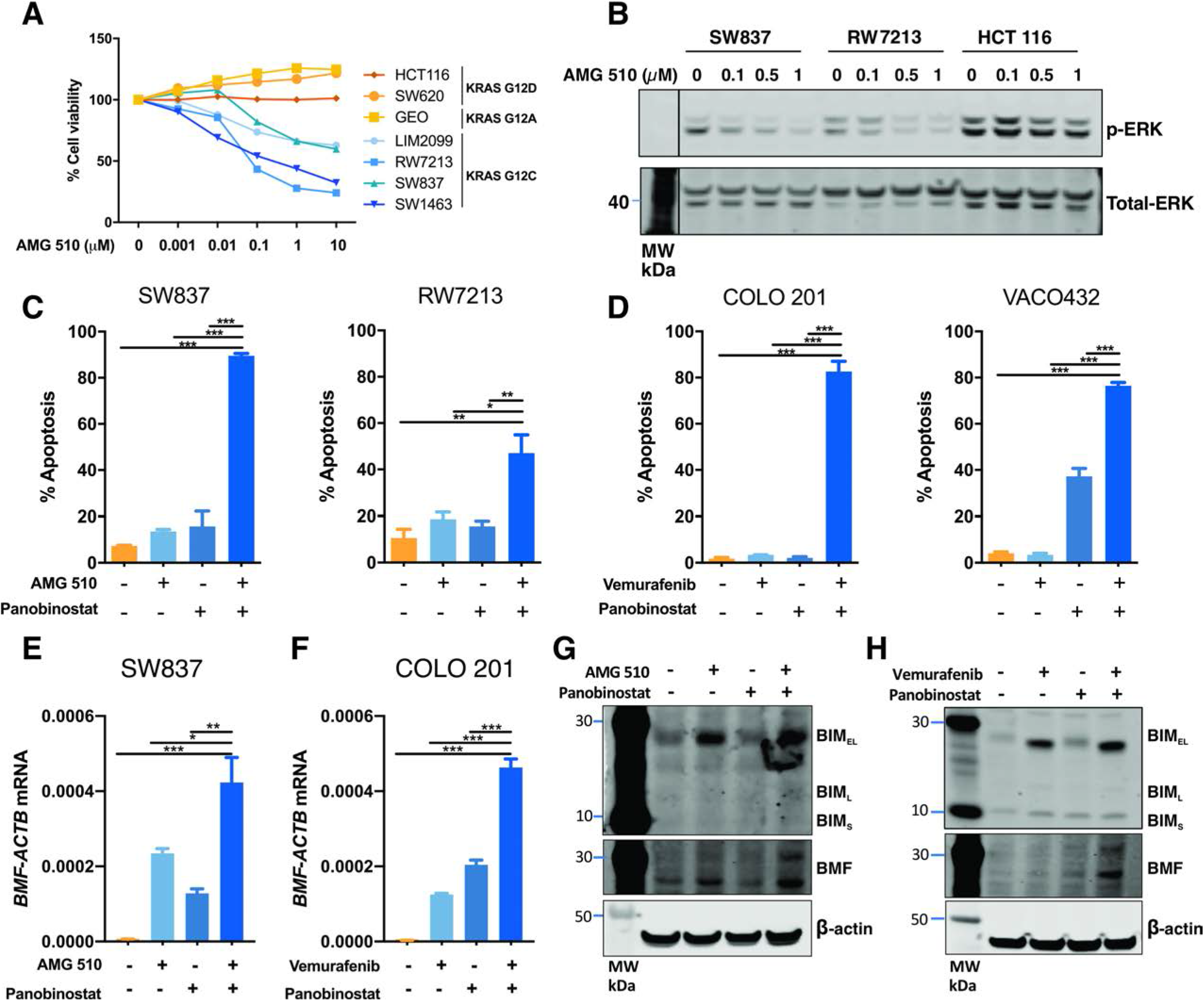
Combined KRAS/HDAC inhibition and BRAF/HDAC inhibition induces apoptosis in CRC cells. (**A**) Effect of AMG 510 on cell viability in CRC cells. *KRAS*^*G12C*^ mutant (LIM2099, RW7213, SW837 and SW1463), *KRAS*^*G12D*^ mutant (HCT 116 and SW620), and *KRAS*^*G12A*^ mutant (GEO) CRC cells were treated with escalating doses of AMG 510 for 72 hours, and cell viability assessed by MTS assay. Values shown are mean ± SEM from a single experiment performed in technical triplicate. (**B**) SW837, RW7213 and HCT 116 cells treated with AMG 510 for 1 hour and p-ERK and total ERK levels determined by western blot analysis. Apoptosis analysis as determined by propidium iodide staining and FACS analysis in (**C**) SW837 and RW7213 cells were treated AMG 510 (0.1 μM) and panobinostat (25 nM) alone and in combination for 72 and (**D**) COLO 201 and VACO432 cells treated with vemurafenib (2.5 μM) and panobinostat (25 nM) for 72 hours. Values shown are mean ± SEM from a single representative experiment performed in technical triplicate. Similar results were obtained in two additional independent experiments. One-Way ANOVA, with Tukey’s multiple comparison testing; *(*p*≤0.05), *** (*p*≤0.001) and ns=not significant. q-RT-PCR analysis of *BMF* mRNA expression in (**E**) SW837 cells treated with AMG 510 (0.1 μM) and panobinostat (25 nM) and (**F**) COLO 201 cells treated with vemurafenib (2.5 μM) and panobinostat (25 nM) for 12 hours. Values shown are mean ± SEM from a single representative experiment performed in technical triplicate; *(*p*≤0.05), **(*p*≤0.01), *** (*p*≤0.001). Western blot analysis for BIM and BMF expression in (**G**) SW837 cells treated with AMG 510 and panobinostat and (**H**) COLO 201 cells treated with vemurafenib and panobinostat for 12 hours. β-tubulin was used as a loading control.

Finally, we assessed the effect of combining panobinostat with the BRAF^V600^ inhibitor vemurafenib in *BRAF*^*V600E*^ cell lines. As observed for the other MAPK genotype-tailored combinations, BRAF^V600^/HDAC inhibition significantly enhanced apoptosis compared to either agent alone (**Figure 6D**), and significantly increased *BMF* mRNA and BIM protein expression (**Figure 6F, H**).

## Discussion

Herein, we demonstrate that combined inhibition of ERK/MAPK signalling, and histone deacetylases, synergistically induces apoptosis across multiple CRC cell lines. Notably, this effect could be generically induced across CRC cell lines by combining a MEK inhibitor with an HDAC inhibitor, or it could be selectively tailored to the ERK/MAPK genotype of CRC cells by combining EGFR, KRAS^G12C^ or BRAF inhibitors with an HDAC inhibitor, in *RAS/BRAF*^WT^, *KRAS*^*G12C*^ or *BRAF^V600^* CRC cell lines, respectively. The latter approach may provide greater tumour-specific targeting and a better therapeutic window compared to MEK inhibition where on-target toxicities in normal cells may reduce clinical activity [34].

The robust pro-apoptotic activity of MEK/HDAC inhibition was also observed in PDTO’s generated from four CRC patients, three of which were derived from metastatic sites. The combination also effectively inhibited colonic adenoma growth in an *Apc* mutant mouse model, illustrating the potential for this regimen to be effective at multiple stages in the adenoma-carcinoma sequence.

Mechanistically, we demonstrate that the synergistic activity of the combination is largely driven by the additive induction of the BH-3 only proteins, BMF and BIM. BIM has a strong binding affinity to all BCL-2 proteins (BCL-2, BCL-X_L_, MCL-1, BCL-W and BFL-1), whereas BMF primarily binds to BCL-X_L_, BCL-W and BCL-2, and only weakly binds to MCL-1 and BFL-1 [35]. As CRCs express high levels of BCL-X_L_ [36], we postulate that the combinatorial induction of both BIM and BMF may be sufficient to bind and inhibit the high levels of these pro-survival factors, enabling BIM to bind and activate BAX/BAK or to inhibit other pro-survival proteins and trigger apoptosis.

Despite previous publications demonstrating *BIM* mRNA induction following MAPKi or HDACi treatment [24, 32], we did not observe this to be the case in all cell lines in this study. Comparatively, we observed a consistent increase in BIM protein expression, particularly with trametinib treatment. BIM is expressed as three isoforms (BIM_EL_, BIM_L_ and BIM_S_) of which BIM_EL_ has been previously shown to be regulated by MAPK signalling [6–8, 33]. Specifically, ERK-mediated phosphorylation of BIM_EL_ at serine-69 promotes its ubiquitination and proteasomal degradation [6, 8], while BIM_EL_ phosphorylation has also been shown to induce its rapid dissociation from the BCL-2 proteins, BCL-X_L_ and MCL-1 [33]. Consistent with these reports, trametinib-mediated inhibition of MEK/ERK signalling induced the strongest increase in protein expression of the BIM_EL_ isoform and induced a decrease in the band size of BIM_EL_, consistent with a loss of phosphorylation event.

In addition, consistent induction of *BMF* mRNA was observed following both MAPK pathway and HDAC inhibition. Notably, the BMF promoter harbours multiple SP1/SP3 binding sites and localisation of SP3 to the BMF promoter has been demonstrated in ChIP studies [21, 23]. HDACi are known to preferentially regulate SP1/SP3 target genes, either through inhibition of HDAC enzymes which are recruited by these factors, or by direct acetylation of SP1 or SP3 [37]. MAPK signalling has also been shown to modulate SP1 activity through direct ERK-mediated phosphorylation of SP1 at Thr-453 and Thr-739 [38], raising the potential that MAPK pathway and HDAC inhibition converge to regulate BMF expression by modulating the activity of SP1/SP3 transcription factors.

Our finding that combined MAPK signalling/HDAC inhibition enhances cell death in CRC cells is reminiscent of studies in other tumour types, although several different mechanisms have been proposed. In triple negative and inflammatory breast cancer, combined MEK/HDAC inhibition enhanced apoptosis by increasing expression of the pro-apoptotic factor NOXA and subsequent inhibition of MCL-1 [39]. Notably, NOXA (*PMAIP1*) expression was not consistently altered in our analysis of the intrinsic apoptotic pathway across the four CRC cell lines, which may reflect differences in tumour type. Several studies have also demonstrated increased cell killing by combined MAPK/HDAC inhibition in melanoma cells, with multiple mechanisms implicated including attenuation of PI3K/AKT signalling and YAP activation [40], suppression of ELK, suppression of the homologous DNA repair and non-homologous end joining pathways [41], and increased ROS production [42]. Collectively, these findings suggest this drug combination may have broad anti-tumour activity across multiple tumour types.

In summary, our finding that combining MAPK pathway inhibitors with HDAC inhibitors can synergistically induce apoptosis in multiple pre-clinical models of colorectal cancer, suggests this concept is worthy of clinical investigation, and we identify strategies in which this can be achieved in a highly targeted manner by tailoring treatment combinations to the specific MAPK genotype of CRCs.

## Author Contributions

L.J.J, I.Y.L, M.P, K.L.S, T.T, G.I, I.N, N.V, J.W.T, R.N, Z.A, S.A, J.M.M: Conducted, analysed and interpreted experiments

L.J.J, A.D, N.C.T, M.E, J.D, O.M.S, P.G, M.P, W.D.F, E.F.L, J.M.M: Conceived, designed, interpreted experiments and/or supervised parts of the study

## Acknowledgements

This project was supported by the Tour de Cure Senior Research grant (17-ONJC-RS-05) and the Operational Infrastructure Support Program, Victorian Government, Australia. LJJ was supported by La Trobe University Australian Postgraduate Awards. GO, IN and ZA was supported by La Trove University Postgraduate Research Scholarship. The patient derived tumour organoid study was supported by the Stafford Fox Medical Research Foundation. JMM (1046092) and OMS (1136119) were supported by National Health and Medical Research Council (NHMRC) Senior Research Fellowships.

---

**Supplementary Figure 1. MEK/HDACi-induced apoptosis is caspase-dependent.** HT29 cells were treated with trametinib (10 nM) and panobinostat (25 nM) alone and in combination in the presence or absence of the pan-caspase inhibitor Q-VD-OPh (10 μM) for 72 hours, and apoptosis determined by propidium iodide staining and FACS analysis. Values shown are mean ± SEM from a single experiment performed in technical triplicate. Similar results were obtained in a second independent experiment. One-Way ANOVA, with Tukey’s multiple comparison testing; *** (*p*≤0.001).

**Supplementary Figure 2. MEK/HDAC inhibition induces apoptosis in CRC cells.** COLO 201 cells were treated with trametinib (10 nM) alone and in combination with valproic acid (2.5 mM), romidepsin (10 nM) or mocetinostat (2.5 μM) for 72 hours and apoptosis induction determined by propidium iodide staining and FACS analysis. Values shown are mean ± SEM from a single experiment performed in technical triplicate. Similar results were obtained in two additional independent experiments. One-Way ANOVA, with Tukey’s multiple comparison testing; *** (*p*≤0.001).

**Supplementary Figure 3.** Representative immunostaining of p-ERK in colonic adenomas and adjacent normal epithelium from *Apc*^*lox/lox*^;*Cdx-Cre*^*ERT2*^ mice. Scale bar = 300 μM

**Supplementary Figure 4. Effect of MEK/HDAC and EGFR/HDAC inhibition on body weight.** Relative mouse weight change compared from commencement of treatment in (**A**) *Apc*^*lox/lox*^;*Cdx-Cre*^*ERT2*^ mice treated with vehicle control (n=5), trametinib (n=4), panobinostat (n=5) and combination (n=8) treated mice and (**B**) BALB/*c* nude mice harbouring LIM1215 xenografts and treated with vehicle control (n=5), cetuximab (n=5), panobinostat (n=5) or the combination (n=5). Error bars represent mean ± SEM.

**Supplementary Figure 4. MEK/HDAC inhibition alters mRNA expression of intrinsic apoptotic pathway components.** COLO201, HT29, DIFI and HCT 116 cells were treated with trametinib (10 nM) and panobinostat (25 nM) alone and in combination for 12 hours. Changes in expression of the pro-apoptotic genes; *BCL2L11*, *BMF*, *BBC3*, *BID*, *BIK* and *PMAIP1* the apoptotic effectors; *BAK*, *BAX* and *BOK* and the Bcl-2 only genes; *BCL2L1*, *MCL-1*, *BCL2*, *BCL2A1* and *BCL2L2* were determined by q-RT-PCR and presented as fold change over DMSO control.

## References

1. Muzny, D.M., et al., Comprehensive molecular characterization of human colon and rectal cancer. Nature, 2012. 487(7407): p. 330–337.

2. McKay, J.A., et al., Evaluation of the epidermal growth factor receptor (EGFR) in colorectal tumours and lymph node metastases. Eur J Cancer, 2002. 38(17): p. 2258–64.

3. Lavoie, H., J. Gagnon, and M. Therrien, ERK signalling: a master regulator of cell behaviour, life and fate. Nat Rev Mol Cell Biol, 2020. 21(10): p. 607–632.

4. Yang, J.Y., et al., ERK promotes tumorigenesis by inhibiting FOXO3a via MDM2-mediated degradation. Nat Cell Biol, 2008. 10(2): p. 138–48.

5. Lin, L., et al., MEK inhibitors induce apoptosis via FoxO3a-dependent PUMA induction in colorectal cancer cells. Oncogenesis, 2018. 7(9): p. 67.

6. Marani, M., et al., Role of Bim in the survival pathway induced by Raf in epithelial cells. Oncogene, 2004. 23(14): p. 2431–41.

7. Hubner, A., et al., Multisite phosphorylation regulates Bim stability and apoptotic activity. Mol Cell, 2008. 30(4): p. 415–25.

8. Ley, R., et al., Activation of the ERK1/2 signaling pathway promotes phosphorylation and proteasome-dependent degradation of the BH3-only protein, Bim. J Biol Chem, 2003. 278(21): p. 18811–6.

9. Shao, Y. and A.E. Aplin, ERK2 phosphorylation of serine 77 regulates Bmf pro-apoptotic activity. Cell Death Dis, 2012. 3: p. e253.

10. Zha, J., et al., Serine phosphorylation of death agonist BAD in response to survival factor results in binding to 14-3-3 not BCL-X(L). Cell, 1996. 87(4): p. 619–28.

11. Bonni, A., et al., Cell survival promoted by the Ras-MAPK signaling pathway by transcription-dependent and -independent mechanisms. Science, 1999. 286(5443): p. 1358–62.

12. Domina, A.M., et al., MCL1 is phosphorylated in the PEST region and stabilized upon ERK activation in viable cells, and at additional sites with cytotoxic okadaic acid or taxol. Oncogene, 2004. 23(31): p. 5301–15.

13. Sale, M.J., et al., Targeting melanoma's MCL1 bias unleashes the apoptotic potential of BRAF and ERK1/2 pathway inhibitors. Nat Commun, 2019. 10(1): p. 5167.

14. Nangia, V., et al., Exploiting MCL1 Dependency with Combination MEK + MCL1 Inhibitors Leads to Induction of Apoptosis and Tumor Regression in KRAS-Mutant Non-Small Cell Lung Cancer. Cancer Discov, 2018. 8(12): p. 1598–1613.

15. Kawakami, H., et al., Mutant BRAF Upregulates MCL-1 to Confer Apoptosis Resistance that Is Reversed by MCL-1 Antagonism and Cobimetinib in Colorectal Cancer. Mol Cancer Ther, 2016. 15(12): p. 3015–3027.

16. Jhawer, M., et al., PIK3CA mutation/PTEN expression status predicts response of colon cancer cells to the epidermal growth factor receptor inhibitor cetuximab. Cancer Research, 2008. 68(6): p. 1953–1961.

17. Mauri, G., et al., Retreatment with anti-EGFR monoclonal antibodies in metastatic colorectal cancer: Systematic review of different strategies. Cancer Treat Rev, 2019. 73: p. 41–53.

18. Siravegna, G., et al., Clonal evolution and resistance to EGFR blockade in the blood of colorectal cancer patients. Nat Med, 2015. 21(7): p. 827.

19. West, A.C. and R.W. Johnstone, New and emerging HDAC inhibitors for cancer treatment. J Clin Invest, 2014. 124(1): p. 30–9.

20. Bolden, J.E., et al., HDAC inhibitors induce tumor-cell-selective pro-apoptotic transcriptional responses. Cell Death Dis, 2013. 4: p. e519.

21. Kang, Y., et al., HDAC8 and STAT3 repress BMF gene activity in colon cancer cells. Cell Death Dis, 2014. 5: p. e1476.

22. Wiegmans, A.P., et al., Deciphering the molecular events necessary for synergistic tumor cell apoptosis mediated by the histone deacetylase inhibitor vorinostat and the BH3 mimetic ABT-737. Cancer Res, 2011. 71(10): p. 3603–15.

23. Zhang, Y., et al., Bmf is a possible mediator in histone deacetylase inhibitors FK228 and CBHA-induced apoptosis. Cell Death Differ, 2006. 13(1): p. 129–40.

24. Zhao, Y., et al., Inhibitors of histone deacetylases target the Rb-E2F1 pathway for apoptosis induction through activation of proapoptotic protein Bim. Proc Natl Acad Sci U S A, 2005. 102(44): p. 16090–5.

25. Rahmani, M., et al., Coadministration of histone deacetylase inhibitors and perifosine synergistically induces apoptosis in human leukemia cells through Akt and ERK1/2 inactivation and the generation of ceramide and reactive oxygen species. Cancer Res, 2005. 65(6): p. 2422–32.

26. Chueh, A.C., et al., ATF3 Repression of BCL-XL Determines Apoptotic Sensitivity to HDAC Inhibitors across Tumor Types. Clin Cancer Res, 2017. 23(18): p. 5573–5584.

27. Duan, H., C.A. Heckman, and L.M. Boxer, Histone deacetylase inhibitors down-regulate bcl-2 expression and induce apoptosis in t(14;18) lymphomas. Mol Cell Biol, 2005. 25(5): p. 1608–19.

28. Mouradov, D., et al., Colorectal cancer cell lines are representative models of the main molecular subtypes of primary cancer. Cancer Res, 2014. 74(12): p. 3238–47.

29. Mariadason, J.M., et al., Gene expression profiling-based prediction of response of colon carcinoma cells to 5-fluorouracil and camptothecin. Cancer Res, 2003. 63(24): p. 8791–812.

30. Hinoi, T., et al., Mouse model of colonic adenoma-carcinoma progression based on somatic Apc inactivation. Cancer Res, 2007. 67(20): p. 9721–30.

31. Aubrey, B.J., et al., An inducible lentiviral guide RNA platform enables the identification of tumor-essential genes and tumor-promoting mutations in vivo. Cell Rep, 2015. 10(8): p. 1422–32.

32. Cragg, M.S., et al., Gefitinib-induced killing of NSCLC cell lines expressing mutant EGFR requires BIM and can be enhanced by BH3 mimetics. PLoS Med, 2007. 4(10): p. 1681–89; discussion 1690.

33. Ewings, K.E., et al., ERK1/2-dependent phosphorylation of BimEL promotes its rapid dissociation from Mcl-1 and Bcl-xL. EMBO J, 2007. 26(12): p. 2856–67.

34. Caunt, C.J., et al., MEK1 and MEK2 inhibitors and cancer therapy: the long and winding road. Nat Rev Cancer, 2015. 15(10): p. 577–92.

35. Chen, L., et al., Differential targeting of prosurvival Bcl-2 proteins by their BH3-only ligands allows complementary apoptotic function. Mol Cell, 2005. 17(3): p. 393–403.

36. Krajewska, M., et al., Elevated expression of Bcl-X and reduced Bak in primary colorectal adenocarcinomas. Cancer Res, 1996. 56(10): p. 2422–7.

37. Chueh, A.C., et al., Mechanisms of Histone Deacetylase Inhibitor-Regulated Gene Expression in Cancer Cells. Antioxid Redox Signal, 2015. 23(1): p. 66–84.

38. Milanini-Mongiat, J., J. Pouyssegur, and G. Pages, Identification of two Sp1 phosphorylation sites for p42/p44 mitogen-activated protein kinases: their implication in vascular endothelial growth factor gene transcription. J Biol Chem, 2002. 277(23): p. 20631–9.

39. Torres-Adorno, A.M., et al., Histone Deacetylase Inhibitor Enhances the Efficacy of MEK Inhibitor through NOXA-Mediated MCL1 Degradation in Triple-Negative and Inflammatory Breast Cancer. Clin Cancer Res, 2017. 23(16): p. 4780–4792.

40. Faiao-Flores, F., et al., HDAC Inhibition Enhances the In Vivo Efficacy of MEK Inhibitor Therapy in Uveal Melanoma. Clin Cancer Res, 2019. 25(18): p. 5686–5701.

41. Maertens, O., et al., MAPK Pathway Suppression Unmasks Latent DNA Repair Defects and Confers a Chemical Synthetic Vulnerability in BRAF-, NRAS-, and NF1-Mutant Melanomas. Cancer Discov, 2019. 9(4): p. 526–545.

42. Wang, L., et al., An Acquired Vulnerability of Drug-Resistant Melanoma with Therapeutic Potential. Cell, 2018. 173(6): p. 1413–1425 e14.

